# Structure and dynamics of secondary and mature rainforests: insights from South Asian long-term monitoring plots

**DOI:** 10.1101/2023.12.24.573243

**Authors:** Akhil Murali, Srinivasan Kasinathan, Kshama Bhat, Jayashree Ratnam, Mahesh Sankaran, Divya Mudappa, T. R. Shankar Raman, Anand M. Osuri

**Author notes:** Corresponding author: Akhil Murali, Nature Conservation Foundation, 1311, 12th A Main, Vijayanagar 1st Stage, Mysuru 570017, Karnataka, India.

## Abstract

We assessed tree community and carbon dynamics over 5 years in two 1-ha long-term ecosystem monitoring plots, one each in mature tropical rainforests (MR) and 10-year post-agroforestry secondary rainforests (SR) in India’s Western Ghats mountains. Both plots were established in 2017 and monitored annually six times. We expected (1) higher tree diversity, distinct species composition, and greater carbon stock in MR; (2) higher carbon sequestration rates in SR; and (3) carbon dynamics shaped by growth and mortality in SR and MR, respectively. The SR plot had fewer species (67 vs. 84), stored substantially less carbon (76 vs. 193 Mg), and comprised a distinct community with fewer late-successional species than MR. SR gained 5.8 Mg carbon, due to tree growth exceeding losses from mortality, while in MR mortality exceeded growth and recruitment resulting in a 3.3 Mg carbon decline over five years. While MR had higher tree diversity, carbon stocks and relatively intact composition, the high rates of biodiversity and carbon accrual in SR highlight the conservation and climate significance of post-agroforestry secondary forests. Moderate carbon losses noted here in MR, as in other mature South Asian tropical forests, is a cause for concern under ongoing climate change.

## Introduction

Tropical forests constitute about 50% of global forest area, hold 55% of total terrestrial carbon (Pan *et al*. 2011), and account for nearly two-thirds of global biodiversity (Artaxo *et al*. 2022). These forests are critical to regulating atmospheric CO_2_ and global climate, sustaining biodiversity, and maintaining global ecological resilience and productivity. Increased human activity has reduced the integrity of tropical forests worldwide, including in tropical Asia, where over half the countries have already witnessed significant forest depletion exceeding 70% (Grantham *et al*. 2020; Laurance 2007).

Secondary forests regrowing after various types of past disturbances now dominate many tropical forest landscapes (Corlett 1995). Although not comparable to mature forests in many key ecological attributes (López-Bedoya *et al*. 2022), secondary forests are likely to play important roles in biodiversity conservation and the maintenance of ecosystem functions in the coming decades (Chazdon *et al*. 2009, 2016; Dent and Wright 2009). Tropical secondary forests have emerged as significant terrestrial carbon sinks due to their wide extent and higher rates of carbon fixation when compared to primary forests (Heinrich *et al*. 2023; Martin *et al*. 2013; Zuidema and Jakovac 2023). However, barring a few noteworthy recent contributions (Meister *et al*. 2012), empirical long-term research and monitoring of vegetation and carbon dynamics in tropical secondary forests is quite scarce as research attention has mostly focused on mature tropical forests (Davies *et al*. 2021; Malhi *et al*. 2021). This knowledge gap is particularly acute in regions such as South Asia. Long-term data and insights from these underrepresented regions can refine our understanding of tropical biodiversity and carbon cycling and increase the accuracy of global carbon budgets.

Mature and secondary tropical forests typically differ in structure, composition, function, and dynamics of tree growth, with the magnitude of change varying with age of the site and intensity of past disturbance. Previous studies have documented lower tree species diversity, basal area and biomass in secondary forests compared to mature forests (Borah *et al*. 2019; Brearley *et al*. 2004; Gatti *et al*. 2015), with the extent of these differences closely linked to the specific forest community and its stage of development. Tropical secondary forests tend to be dominated by early-successional light-demanding plant species that are typically fast growing, soft-wooded and short-lived (Bazzaz and Pickett 1980; Lugo and Brown 1992; van Breugel *et al*. 2006).

Long-term studies also reveal differences in temporal dynamics between mature and secondary forests, especially the dynamics of carbon sequestration and the impact of climate change on the successional trends. Rates of carbon sequestration in young recovering secondary forests (<20 years) can be as much as 20 times greater than in mature forests, although their sink potential can vary considerably depending on age, geography, climate, edaphic factors and disturbance regimes (Chave *et al*. 2020; Chazdon *et al*. 2016; Heinrich *et al*. 2023; Matos *et al*. 2020). Ultimately, the carbon balance of forest ecosystems, and whether they function as carbon sinks or sources, hinges on the contrast between net primary productivity and the carbon lost due to tree mortality. Carbon losses due to tree mortality may be balanced by gains due to primary productivity in mature tropical forests. However, in secondary forests, the combination of higher growth and mortality rates may result in a different pattern (Chave *et al*. 2008; Phillips *et al*. 2016). Understanding and quantifying the working of these processes in both mature and secondary forests across diverse geographies is essential, as it not only provides a more accurate quantification of the global carbon sequestration potential of secondary forests, but also informs sustainable land-use decisions and advances our understanding of ecosystem responses to environmental change.

Here, we compare stand structure and short-term dynamics over a 6-year period (2017 - 22) in two 1 ha long-term forest plots in mature and secondary tropical wet evergreen forests of the Anamalai Hills, Western Ghats, India. We expected that as compared to the mature forest, secondary forest would have lower biomass, carbon, and tree diversity, and would vary substantially in species composition. Further, we expected that the regrowing secondary forest would be accumulating carbon faster, while the mature forest would remain relatively stable. Finally, we expected that growth rates would drive carbon dynamics in the secondary forest, while mortality would play a greater role in the primary forest.

## Study Area

Our study focused on the tropical rainforests of the Valparai Plateau and the Anamalai Tiger Reserve, located in the Anamalai Hills, an important conservation area in the southern Western Ghats, India (Fig. 1). The Western Ghats, along with Sri Lanka, has been recognized as a global biodiversity hotspot with 48% of the plant species and 30% of the vertebrates in this region listed as endemic (Kumar *et al*. 2004; Myers *et al*. 2000). The Anamalai Tiger Reserve (ATR, core zone: 958 km^2^, 10.216°N, 76.816°E – 10.566°N, 77.416°E), ranging in elevation between 400 m and 2600 m, contains diverse vegetation types from drier forests in the eastern foothills lying in the rain-shadow to tropical wet evergreen forest types at middle and higher elevations, including on the Valparai Plateau. The Valparai Plateau (220 km^2^, 10.25°N, 76.866°E – 10.366°N, 76.983°E), lying between 700 m and 1500 m, and surrounding parts of ATR receive about 2400 mm of rainfall annually, about two-thirds of which falls during the south-west monsoon between June and August. The rainforest vegetation between 700 m – 1400 m above sea level is characterized as mid-elevation tropical wet evergreen rainforest of the *Cullenia exarillata–Mesua ferrea–Palaquium ellipticum* type (Pascal 1988; Pascal *et al*. 2004).

**Figure 1:**
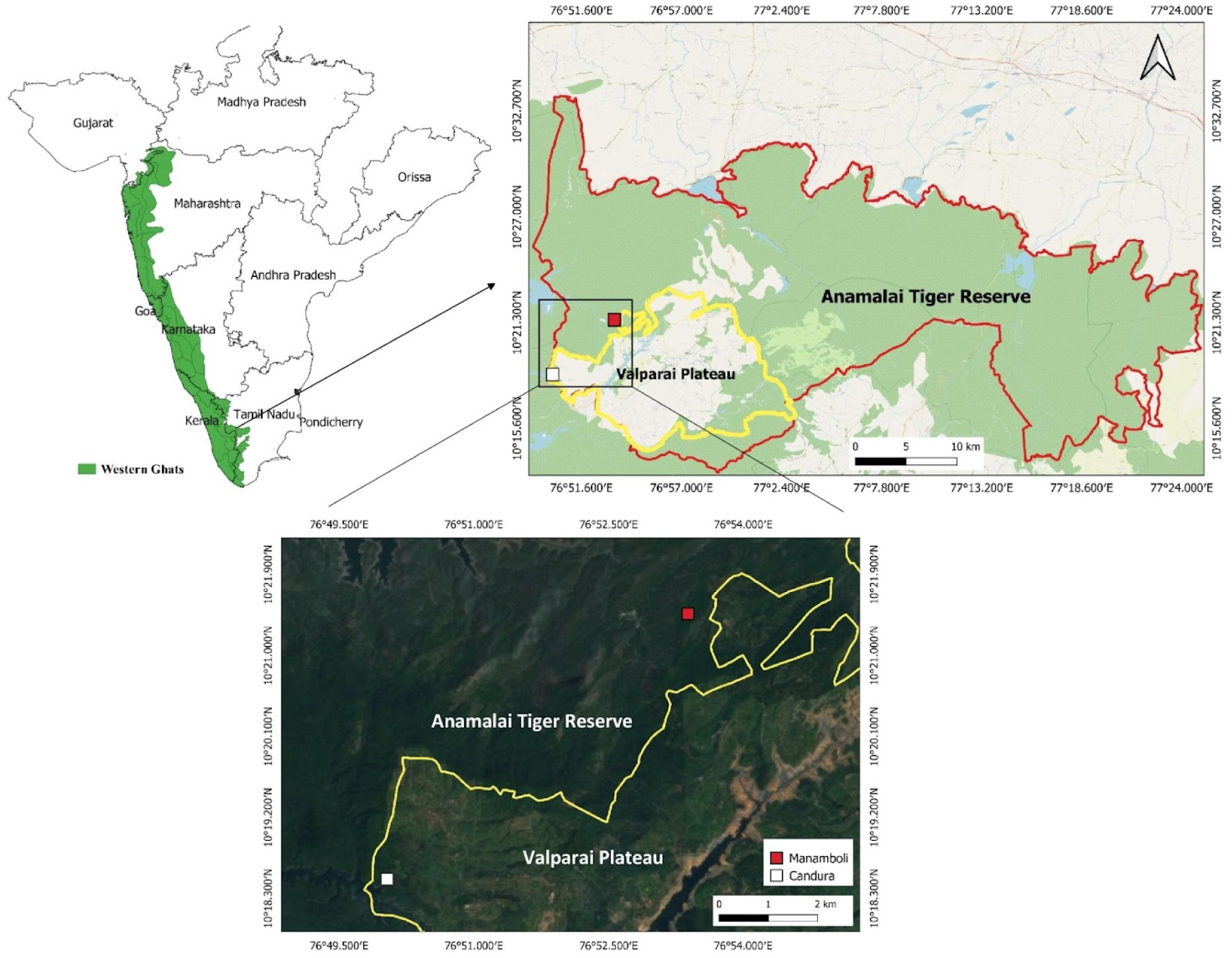
Map showing location of the Anamalai Tiger Reserve and Valparai Plateau in the southern Western Ghats and locations of the two long-term ecological monitoring plots.

The Valparai Plateau is dominated by tea and coffee plantations, first established between 1890 and about 1920 by clearing tropical rainforests. Around 1,100 ha of rainforests remain on the plateau scattered as 50 rainforest fragments ranging from 0.5 ha to 300 ha in size, many of which are degraded due to past deforestation, fragmentation, or conversion to plantations followed by abandonment (Mudappa and Raman 2007; Muthuramkumar *et al*. 2006).

### Selection of Forest Plots

The present study is based on data from two 1 ha forest plots established in 2017—one in mature tropical rainforest and one in secondary forest—both located in the mid-elevation tropical wet evergreen rainforest zone on the western side of the Valparai Plateau (Fig. 1). The mature forest plot is located at Manamboli (10.357748°N, 76.889747°E; 825 m asl), a relatively undisturbed 200 ha rainforest tract within the core area of ATR, protected from logging or other major disturbances since 1979 when the protected area was established. The mature forest is relatively intact and dominated by tall, wet evergreen forest vegetation with characteristic mid-elevation tree species such as *Cullenia exarillata*, *Palaquium ellipticum*, and *Mesua ferrea*, besides species such as *Paracroton pendulus*, *Reinwardtiodendron anamalaiense,* and the dipterocarp *Vateria indica* (Muthuramkumar *et al*. 2006) at lower elevations. The Manamboli rainforest faces some disturbance due to access roads passing through and because of robusta coffee (*Coffea canephora*) invasion in a section adjoining a neighbouring plantation (Joshi *et al*. 2009). The location selected for the 1 ha plot avoided the robusta invaded area and was over 100 m away from the main road passing through Manamboli and about 50 m from the edge of a smaller forest road.

The secondary forest plot is located in a 124-ha rainforest remnant adjoining tea plantations on the Valparai Plateau: The Candura rainforest remnant (10.308554°N, 76.833919° E; 875 m asl). The Candura plot lies within Murugalli Estate (Parry Agro Industries Limited, PAIL), on the western edge of the Valparai Plateau. The Candura remnant borders the backwaters of the Lower Sholayar reservoir to the south and has tea plantations to the north and east forming a small enclave within this landscape. Towards the west, Candura abuts extensive contiguous mature forest tracts in Vazhachal Reserved Forest, Kerala. The Candura site has been episodically selectively logged in the 1990s and early 2000s (last episode in 2004). The understorey of the remnant was cleared for cultivation of Vanilla (*Vanilla planifolia*) in the central and southern part in the early 2000s (abandoned in 2007), robusta coffee (*Coffea canephora*) in the northwestern corner (abandoned in early 2000s), and pepper in 21 ha in the northeastern part (established in 2015, abandoned in 2021). These areas were shaded plantations established under scattered native shade trees (such as *Vateria indica, Bombax ceiba*) that remained after logging and supplemented by planted alien shade tree species such as silver oak (*Grevillea robusta*), African musizi (*Maesopsis eminii*), African tulip (*Spathodea campanulata*) and quickstick (*Gliricidia sepium*). Starting from a smaller area in 2007, an area of 103 ha of Candura was set aside for protection and restoration under a 2015 memorandum of understanding between the Nature Conservation Foundation (NCF) and PAIL, to which the abandoned pepper area was added in 2023. The location selected for the 1 ha secondary forest plot was in the center of Candura remnant in secondary forest recovering after past logging and abandonment of vanilla cultivation where no active restoration efforts have been carried out. A small internal access road passes just south of the site.

## Methods

### LEMoN-India Protocol

The two 1 ha plots of 100 m × 100 m dimensions (aligned to N-S and E-W axes) were established in 2017 as part of the Long-term Ecological Monitoring Network – India (LEMoN – India) (Marthews *et al*. 2015). LEMoN forest dynamics plots largely followed the RAINFOR-GEM protocol (Marthews TR *et al*. 2014) with elements of the CTFS protocol (Condit 1998). Each 1 ha plot, sub-divided into 100 continuous sub-plots of 10 m × 10 m, was surveyed and mapped to maximum accuracy using a theodolite in the field, with grid corners permanently staked. All woody plant individuals with girth at breast height (GBH, at 1.3 m) ≥10 cm were tagged with numbered aluminum tags and spatially mapped. Plant species were identified using standard floral keys (Gamble and Fischer 1935; Page 2017; Pascal and Ramesh 1987) and updated to current taxonomy (POWO 2023). Stem GBH was measured for all single stemmed individuals. For trees with buttresses, the GBH point of measurement (POM) was taken at 50 cm above the buttresses or at the height where the stem is regular. GBH of all stems ≥10 cm were measured for trees that were multi-stemmed at the 1.3 m POM. The plots were also recensused each year (around November). New saplings that recruited into the ≥10 cm GBH class were identified, mapped, tagged, and added to the monitoring. Stems that appeared to be dead were recorded at each monitoring and those that showed no signs of recovery in subsequent visits were recorded as mortality. The height of all individual stems was measured using a laser rangefinder (Nikon Monarch 7i VR) during plot establishment as the length along the main stem of the tree. Height was recorded for new recruits added on annual basis, and was re-measured for all stems after 5 y had elapsed (i.e., during the 6^th^ annual census) since plot establishment.

Trees with GBH ≥20 cm were fitted with stainless steel dendrometer bands, attached 20 cm above the point of measurement with the change in circumference since installation indicated by the band measured using vernier calipers. Dendrometer band measurements of all stems were measured in each plot on a quarterly basis.

### Analysis

The basal area of multi-stemmed individuals was calculated using the effective diameter of individual plants measured as the square root of the sum of squared diameters of all stems. We then estimated Above Ground Biomass (AGB) of individual stems using the allometric equation developed for tropical forests (Chave *et al*. 2014):

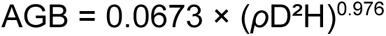

where *ρ* is the specific wood density (gcm^−3^), D is the diameter (cm), H is the tree height (m).

Wood density was sourced from a functional trait data set of trees in the Anamalai hills (Osuri *et al*. 2022) and the global wood density database (Zanne *et al*. 2009).

We evaluated differences in diversity and stand structure between mature and secondary forest by comparing mean annual plot level values of species richness (number of species in 1 ha), density (individuals/ha), basal area (BA, m^2^ha^−1^) and carbon (AGC, Mg/ha) across the six censuses. To convert AGB into AGC we used the following equation: AGC= AGB×0.47 (Hughes *et al*. 1999).

Species composition and relative abundance (based on number of individuals or basal area) were compared across plots using species rank-abundance curves. To evaluate temporal changes in composition, we grouped species into three guilds based on habitat affinity as mature rainforest species, secondary successional species, and introduced (alien) species (Osuri *et al*. 2022).

Stem growth rate was calculated for each tree over the five consecutive annual censuses as the change of diameter per year from the initial to the final year. We converted absolute growth rate to the DBH specific growth rate (Pommerening and Muszta 2016) using the following formula:

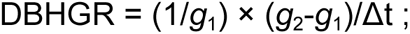

where DBHGR is DBH specific growth rate (cm·cm^−1^·y^−1^), *g*_1_ is initial diameter (cm), *g*_2_ is final diameter (cm), and Δ*t* is the difference in years between two consecutive censuses.

We estimated annual mortality and recruitment as percentages of total stems, based on individuals recorded as dead or added as recruits in annual censuses. We estimated stem net primary productivity (NPP) as the mean annual increase in carbon in live trees (Mg C). To assess the contribution of tree growth, mortality and recruitment to carbon dynamics we partitioned change in C separately into three components averaged across years: increase due to stem recruitment, increase due to stem growth, and loss due to tree mortality.

All analyses were performed with R software (R Core Team 2022). We employed the *tidyverse* package version 1.3.0 (Wickham *et al*. 2019) for handling and organizing our data. Above ground biomass was estimated using the *BIOMASS* R package (Réjou-Méchain *et al*. 2017) and *ggplot2* was used for preparing figures (Wickham *et al*. 2016). Our approach to statistical analysis was to obtain accurate and robust parameter estimates and measures of variability across years, documenting what were evidently clear differences between secondary and mature forests. Instead of statistical significance tests and *P* values, we take a “clean, careful quantitative analysis” approach to produce, understand, and interpret new ecological knowledge (Steel *et al*. 2013). Data and code from the study are available at (Murali *et al*. 2023).

## Results

### Forest structure and species composition

Across the six annual censuses, the mature forest plot averaged fewer (1318) individuals in 34 families but greater diversity at the level of genera (69) and species (84) than the secondary forest plot (1934 individuals, 35 families, 56 genera, and 67 species, Table 1). Despite having fewer stems, the mature forest had a higher average basal area and above ground carbon (Table 1). While the basal area was only 38% higher in mature forest relative to secondary forest (41.4 m^2^ha^−1^ versus 30 m^2^ha^−1^), the above ground carbon stock was nearly three times higher relative to secondary forest (193.47 MgC versus 76.61 MgC, Table 1).

**Table 1:**
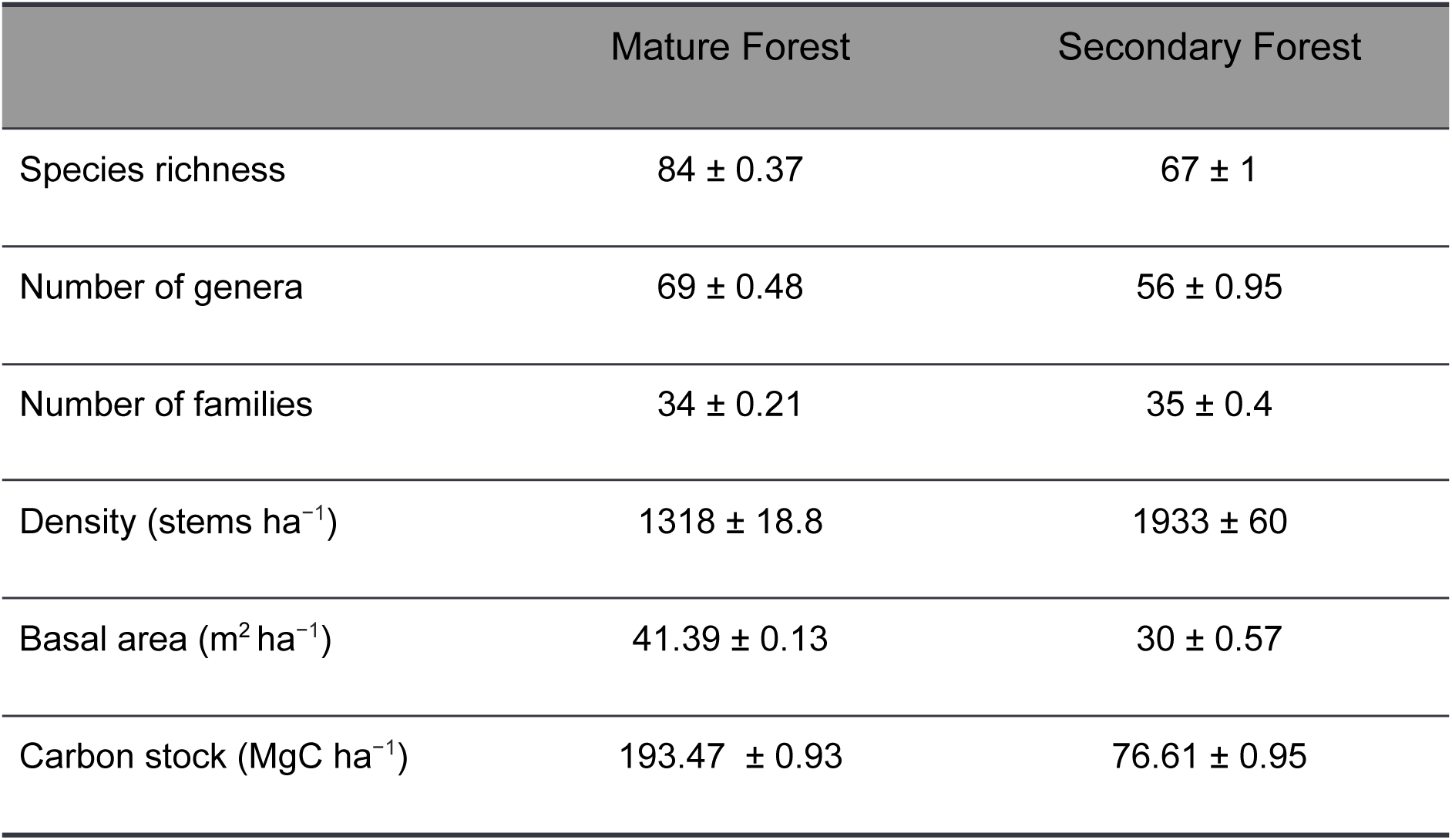
Structural differences between mature and secondary forest plots in mid-elevation tropical wet evergreen forest of the Anamalai Hills (tabled values are means ± SE across six censuses).

Species composition varied distinctly between the two forest plots; in the mature forest, species like *Reinwardtiodendron anamalaiense, Otonephelium stipulaceum,* and *Paracroton pendulus* dominated in terms of stem density, whereas in the secondary forest, *Maesopsis eminii, Strobocalyx arborea,* and *Actinodaphne wightiana* were the dominant species (Fig. 2)*. M. eminii* and *Spathodea campanulata* occurring in the secondary forest plot are known invasive alien species. Invasive alien species were more prevalent (4 species, 15% stems) in secondary forest than mature forest (1 species, 4% stems). In mature forest, 50% of the basal area was contributed by the five most abundant tree species: *Palaquium ellipticum* (8.70 m^2^), *Vateria indica* (5.03 m^2^), *Paracroton pendulus* (2.73 m^2^), *Ficus tsjakela* (2.12 m^2^), and *Reinwardtiodendron anamalaiense* (2.05 m^2^, Fig. 2). In secondary forest, species like *Elaeocarpus tuberculatus (*4.12 m^2^), *Maesopsis eminii* (3.39 m^2^), *Strobocalyx arborea* (3.09 m^2^), *Turpinia malabarica* (2.71 m^2^), and *Vateria indica* (2.19 m^2^) accounted for 50% of the total basal area (Fig. 2).

**Figure 2:**
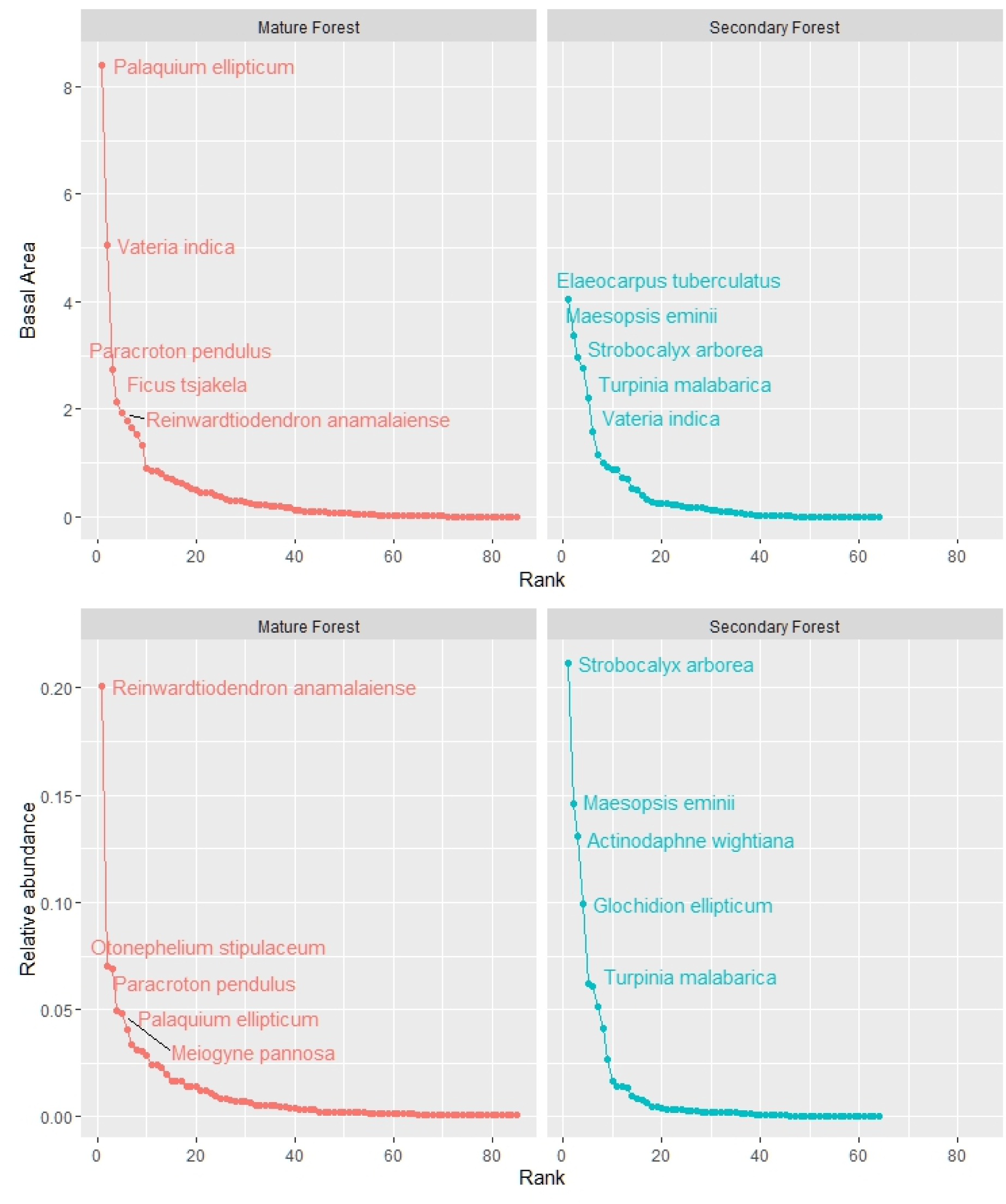
Species rank-abundance curves for mature and secondary forest based on relative abundance of stems (top panel), and basal area (lower panel) based on 2017 data.

### Temporal variation

Over a period of five years (2017–22), temporal changes were more pronounced in the secondary forest (Table 2, Fig. 3). Stem density increased in both plots, with greater total increase (22%), and a mean annual increase of 76 trees ha^−1^ in secondary forest than in the mature forest (11%, 27 trees ha^−1^y^−1^; Table 2). Species richness increased from 64 to 71 species in secondary forest, but remained relatively stable at around 83 species in mature forest. Overall, basal area increased by 12.5% (3.61 m^2^) in the secondary forest, but showed a small decline of 0.6% (0.25 m^2^) in mature forest over the six censuses. Secondary forest accumulated carbon over the five years increasing by 8% from 74.04 MgCha^−1^ to 79.90 MgCha^−1^, with a mean increase of 1.14 MgCha^−1^y^−1^. In contrast, above ground carbon stock in mature forest decreased from 195.16 MgCha^−1^ to 191.81 MgCha^−1^, corresponding to an average drop of 0.67 MgCha^−1^y^−1^ (Table 2).

**Figure 3:**
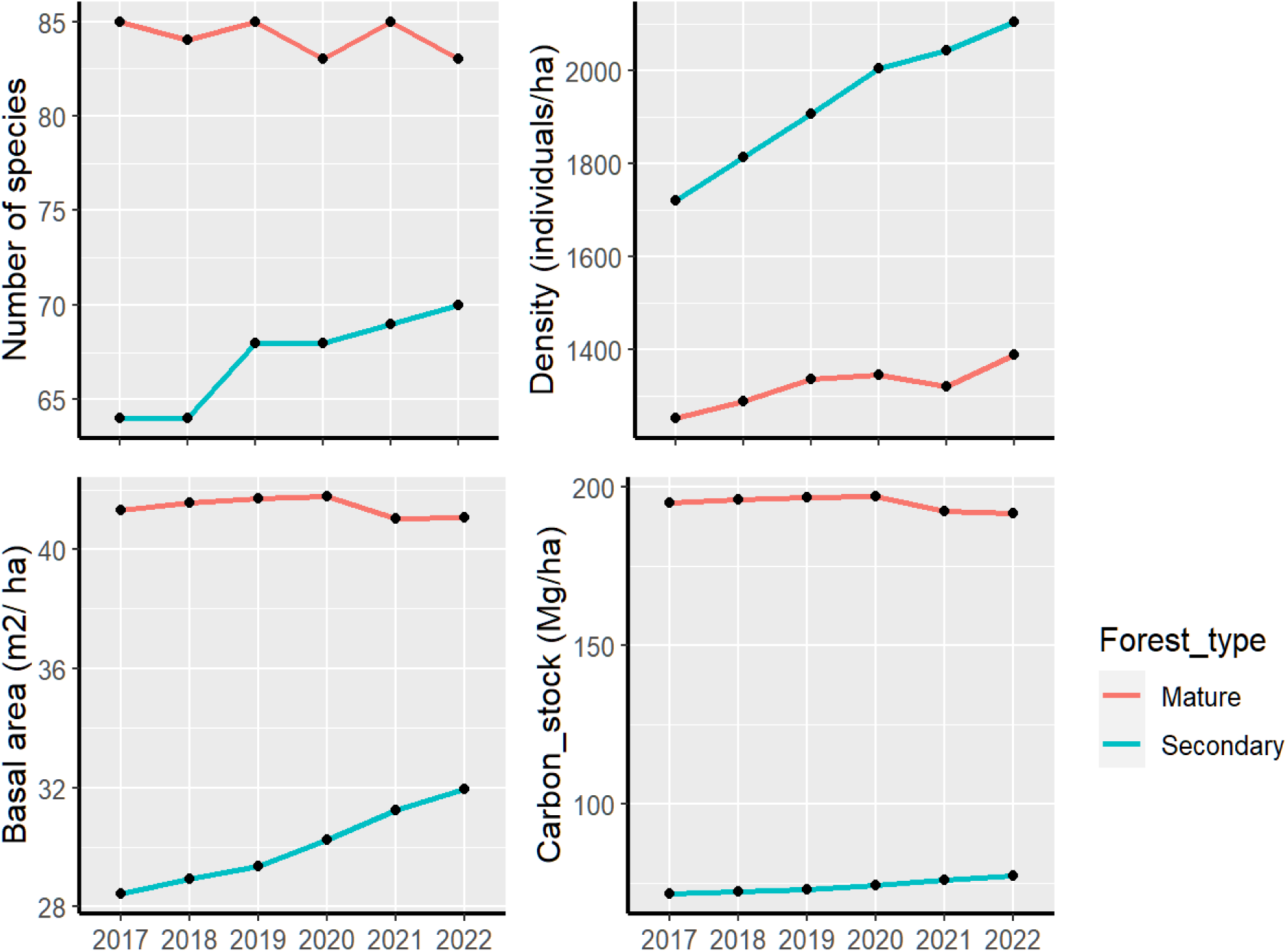
Variation across six years in species richness, density, basal area and above ground carbon stock in mature and secondary forest plots.

**Table 2:** Comparing forest stand structure between the first (2017) and most recent (2022) census in mature and secondary forest (net change is in original units, percentage change in parentheses).

|  | Mature Forest |  |  |  | Secondary Forest |  |  |  |
| --- | --- | --- | --- | --- | --- | --- | --- | --- |
|  | 2017 | 2022 | Net<br>change | Mean<br>annual<br>change<br>(±SE) | 2017 | 2022 | Net<br>change | Mean<br>annual<br>change<br>(±SE) |
| Species richness | 85 | 83 | -2 (2.3%) | -0.40 ±<br>0.81 | 64 | 71 | 7 (10%) | 1.80 ±<br>0.73 |
| Density | 1251 | 1385 | 134 (11%) | 27.60 ±<br>16.19 | 1721 | 2112 | 391<br>(22.31%) | 76.80 ±<br>11.50 |
| Basal area<br>(m<sup>2</sup> ha<sup>-1</sup>) | 41.31 | 41.05 | -0.25<br>(0.61%) | -0.05 ±<br>0.17 | 28.42 | 32.03 | 3.61<br>(12.45%) | 0.71 ±<br>0.11 |
| Carbon stock<br>(MgC ha<sup>-1</sup>) | 195.16 | 191.81 | -3.35<br>(1.71%) | -0.67 ±<br>1.02 | 74.04 | 79.90 | 5.86<br>(8%) | 1.14 ±<br>0.25 |

Over the 5-year span, eight species recruited into the secondary forest plot (*Calophyllum polyanthum, Beilschmiedia* sp.*, Phoebe lanceolata, Actinodaphne tadulingamii, Breynia vitis-idaea, Tabernaemontana* sp.*, Tetrapilus dioicus, Cryptocarya anamalayana*), while one species *Gmelina arborea* was lost and had 5 individuals in the first census. In the mature forest plot, three new species recruited (*Diospyros* sp.*, Diospyros ebenum, Nothopegia* sp.) while five species were lost (*Lasianthus jackianus*, *Ficus exasperata*, *Antiaris toxicaria*,Symplocos sp. and *Strobilanthes* sp.).

Over the 5-year time span, increases in stem density in mature forest was mainly due to small increases in stems of both mature forest and an alien species, *Coffea canephora*. In secondary forest, the number of stems of alien and secondary successional species remained stable and most of the increase in stem density was contributed by mature forest species (Fig. 4).

**Figure 4:**
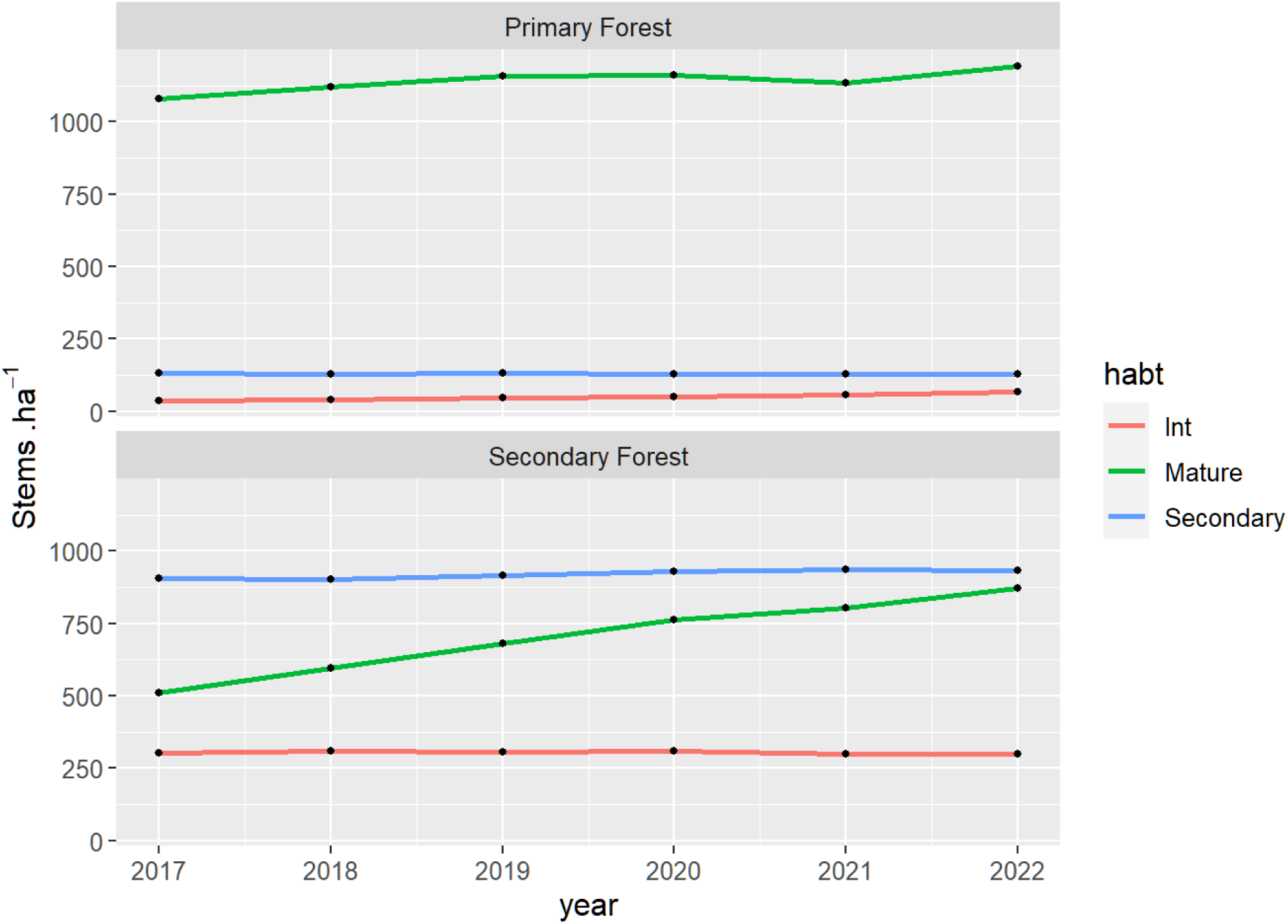
Change in stem density of species belonging to different habitat affinity guilds in secondary and mature forest across the six censuses.

### Growth, recruitment, and mortality

Mortality and recruitment rates were higher in secondary forest than in mature forest (Fig. 5). Annual mortality rates in mature forest ranged between 1% and 2% (except for 4.2% in 2020 due to the high stem mortality of *Strobilanthes* sp., a monocarpic woody shrub). Mortality rates in secondary forest ranged from 2.1% to 3.9%. The secondary forest plot showed higher annual recruitment rates (7% to 8.5%) than mature forest (4.4% to 6.5%). Overall, across both forest types, recruitment rates were higher than mortality rates each year (contributing to net increase in stem count), except for the mature forest plot in 2020.

**Figure 5:**
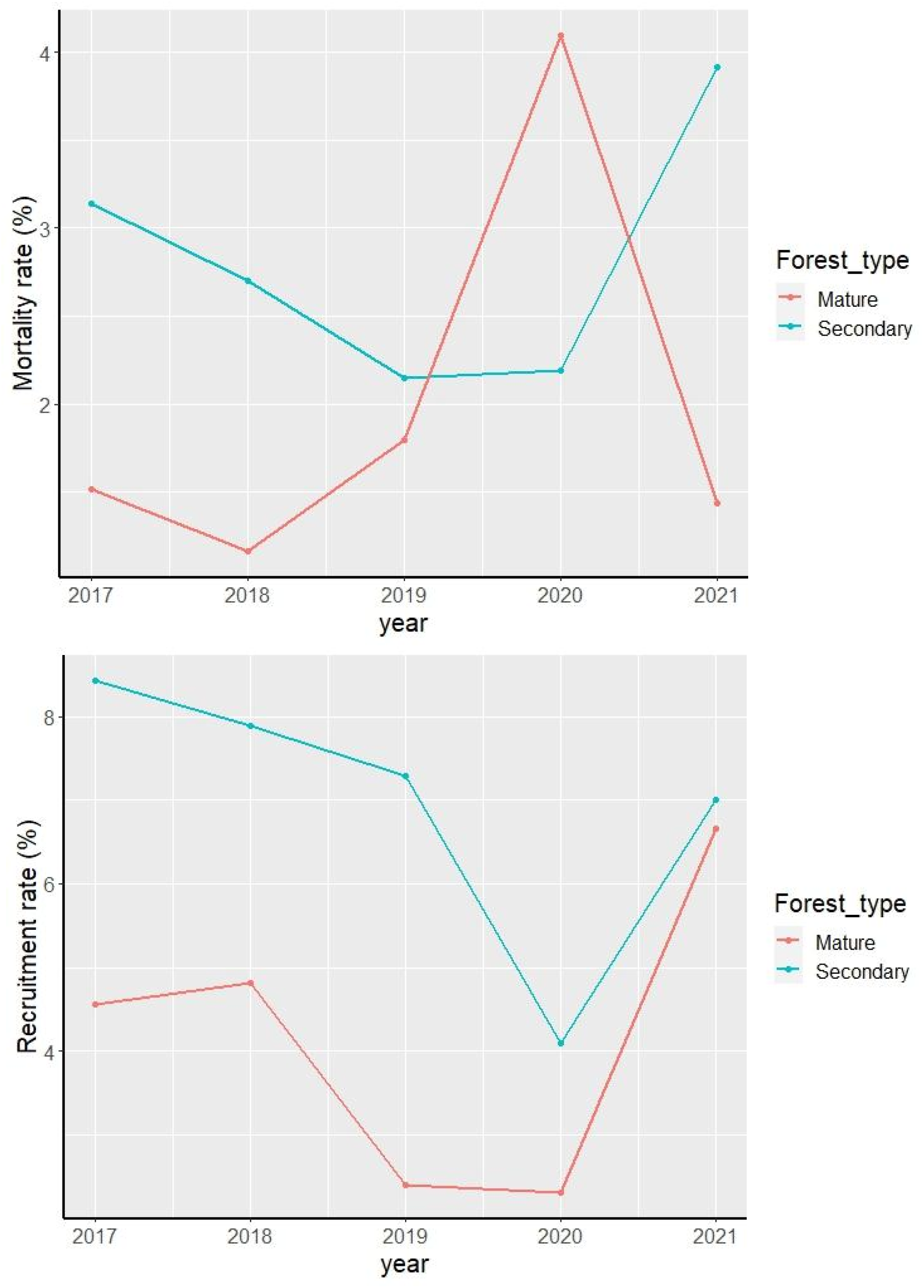
Mortality (top panel) and recruitment (bottom panel) rates in secondary and mature forest across the 5-y time span.

Mean diameter growth rate (DBH specific growth rate), averaged year wise across all stems, was consistently higher in secondary forest ranging between 0.04 cm·cm^−1^y^−1^ and 0.063 cm·cm^−1^y^−1^ than in mature forest where it ranged between 0.004 cm·cm^−1^y^−1^ and 0.026 cm·cm^−1^y^−1^ (Fig. 6).

**Figure 6:**
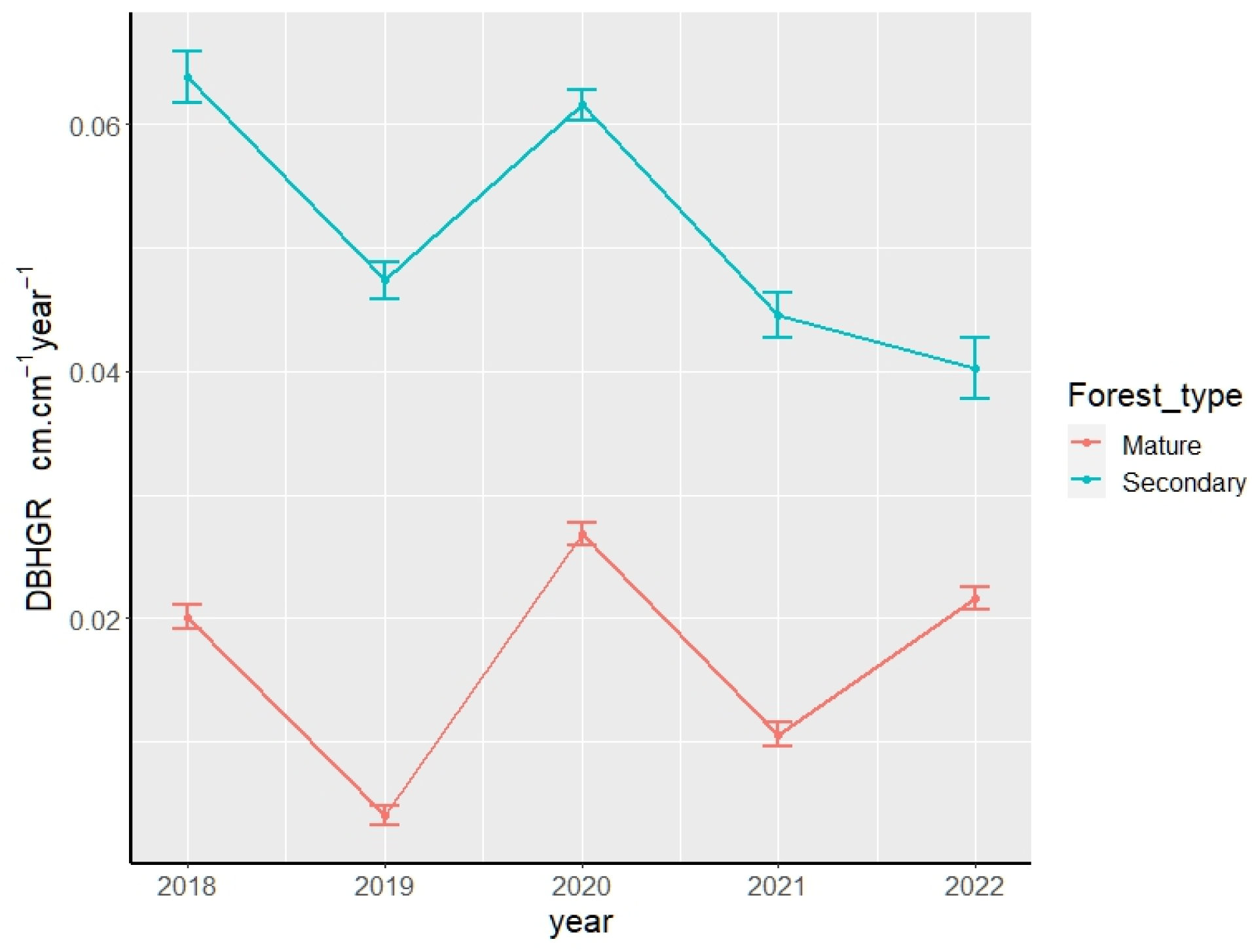
Stem diameter growth measured as diameter at breast height (DBH) specific growth rates in secondary and mature forests (errors bars are standard errors around computed individual tree growth rate).

### Carbon dynamics due to stem growth versus mortality

Change in net stand-level carbon in secondary and mature forests was related to varying contributions of different components of C gain or loss (Fig. 7). Carbon accumulation in secondary forest was mainly contributed by woody stem NPP (2.07 MgCha^−1^y^−1^), with a minor influence of recruits (0.1 MgCha^−1^y^−1^). As C gain was higher than C loss due to stem mortality (1.1 MgCha^−1^y^−1^), secondary forests gained an estimated average 1.14 MgCha^−1^y^−1^. Similarly, woody stem NPP contributed significantly to carbon accumulation in mature forest (1.88 MgCha^−1^y^−1^), while recruits had a negligible influence (0.05 MgCha^−1^y^−1^). However, greater average carbon loss due to tree mortality (2.51 MgCha^−1^y^−1^) led to a reduction in the total stand level carbon in mature forest contrasting with the steady increase in secondary forest.

**Figure 7:**
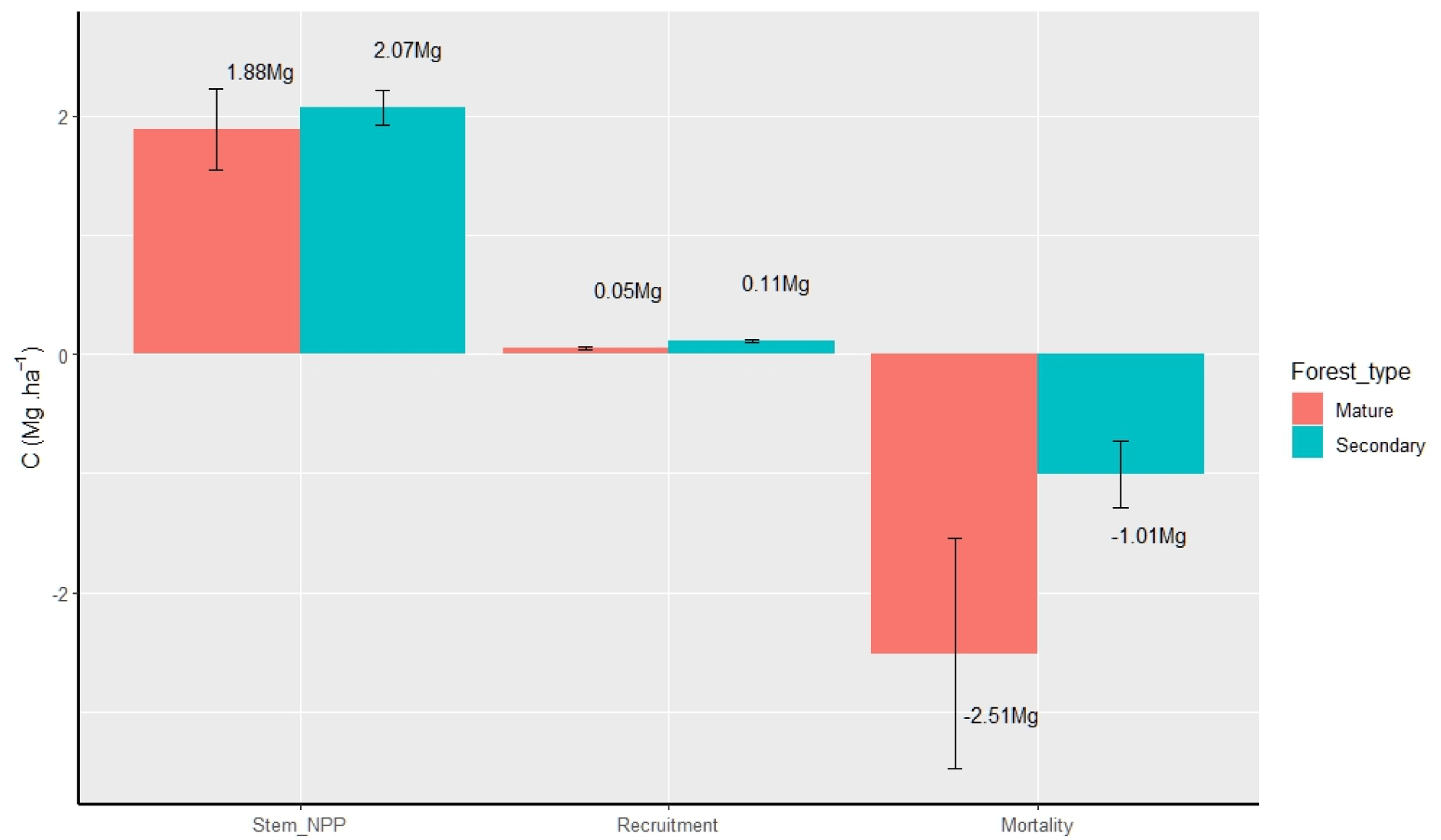
Relative contribution of stem growth, recruitment, and mortality to above ground carbon stocks in secondary and mature forest over the 5-y time span 2017-22 (errors bars are standard errors around computed annual means).

## Discussion

Our assessment of the structure and 5-year dynamics of mature and secondary rainforest plots highlights the distinct and complementary contributions of both types of forests to biodiversity conservation and carbon sequestration. As expected, the mature forest had greater tree diversity, was dominated by shade-tolerant old growth species, and stored nearly three times as much carbon as the secondary forest, which comprised mainly early-successional and non-native species in the adult tree stratum. These are consistent with patterns reported elsewhere across the tropics (Bischoff *et al*. 2005; Brearley *et al*. 2004), reiterating that mature tropical forests hold irreplaceable pools of biodiversity and carbon (Gibson *et al*. 2011; Noon *et al*. 2021).

While the secondary forest did not match the tree diversity, species composition, and carbon stocks of the mature forest, the former was the more dynamic of the two. The secondary forest gained 5.8 MgCha^-1^ over five years, driven largely by tree growth and comparatively low tree mortality. By contrast, the mature forest lost 1.5 times more carbon to mortality than it gained from tree growth, resulting in a decline of 3.3 MgCha^-1^ over the same period. The magnitude and rate of carbon accrual in the secondary forest is noteworthy, considering that active agroforests in India gain an estimated 6.68 MgCha^-1^ over 30 years (Ajit *et al*. 2017; Gopalakrishna *et al*. 2022). It suggests that the potential for climate mitigation in post-agroforestry secondary forests may be higher than previously believed, at least under some conditions, and deserves greater attention in policies and plans for terrestrial carbon sequestration.

In secondary forests regrowing on treeless former land-uses such as open agriculture or pasture, early- to mid-succession is known to be dominated by recruitment of shade-intolerant pioneer species (Chazdon *et al*. 2010; Finegan 1996). By contrast, the post-agroforestry secondary forest at a comparable stage registered significant recruitment of old growth species. Among the recruits are species of particular conservation significance such as *Myristica beddomei* and *Litsea nigrescens* (classified Vulnerable and Endangered, respectively, by the IUCN), alongside numerous other old-growth-associated Western Ghats endemics. The recruitment of old-growth species can also initiate greater long-term carbon capture at this early successional stage because such species grow larger and have more carbon-dense wood than pioneers (Osuri *et al*. 2017; Van Der Sande *et al*. 2016). This early onset of old-growth recruitment could be due to: (a) the retention of some old-growth shade trees in the vanilla and coffee agroforestry plantations prior to abandonment, and (b) the occurrence of mature forests in nearby protected areas. Together, these features have likely enhanced seed arrival and establishment of shade loving old-growth species, both from the native tree species retained in the plantations and the arrival of seeds from nearby mature forests via animal dispersers (Gopal *et al*. 2020; Osuri *et al*. 2017). There is evidence from across the world that carbon stocks in mature tropical forests show distinct regional trends. For example, mature Amazonian forests lost carbon during 1985-2015 whereas carbon in mature African forest remained stable over this period (Hubau *et al*. 2020). There are indications that mature Asian tropical forests are gaining carbon, but this is informed entirely by plots in east and southeast Asia (Suarez *et al*. 2019). South Asia lacks a comparable network of long-term plots for robustly investigating regional trends, but three long term datasets that we are aware of (including this study) indicate recent carbon declines in mature rainforests in the region. While the mature forest plot in our study lost 3.3 MgC over five years, a mature rainforest in the central Western Ghats lost 16.8 Mg biomass (approximately 0.8 MgCha^-1^y^-1^) over a recent 10 year period (Wilson et al. 2023). Stand basal area and aboveground biomass (a strong correlate of carbon storage) decreased over 40 years (basal area from 41.8 m^2^ ha^-1^ to 35.2 m^2^ ha^-1^ or by 15.8%; aboveground biomass from 517.52 Mg ha^-1^ to 430.11 Mg ha^-1^) in a Sri Lankan rainforest (Ediriweera *et al*. 2020, 2023). These three examples suggest that mature South Asian tropical forests could be on a different carbon trajectory from other Asian forests, adding to the reasons for expanding long-term forest monitoring in this historically understudied region.

## Conclusions

Our results underscore the values of protecting relatively intact mature tropical forests for conservation of biodiversity and as reservoirs of carbon. However, the moderate carbon losses observed in mature forests adds to growing evidence of similar trends in other South Asian sites and signal the need for long-term monitoring, especially of tree mortality rates, and incorporating regional estimates into global carbon budgets. Secondary forest recovering after agroforestry use, showing high rates of recovery of biodiversity and carbon accrual (5.8 MgCha^-1^ over five years), can help extend conservation to landscapes outside protected areas and sequester significant amounts of carbon in human-modified tropical landscapes.

## Acknowledgements

We thank the Tamil Nadu Forest Department for research permits and support, especially V. Ganesan, I. Anwardeen, and S. Ramasubramanian, besides a number of DFOs, range officers, foresters, and field staff. For funding and partnership we are grateful to Rohini Nilekani Philanthropies, AMM Murugappa Chettiar Research Centre, Rainmatter Foundation, and Arvind Datar. AMO thanks the Department of Biotechnology (DBT India) for the Ramalingaswami Re-entry Fellowship support (Grant BT/RLF/Re-entry/58/2018). We thank Siddarth Machado, S. K. Chengappa, and H. V. Raghavendra for help with setting up the plot and Mrinalini K. Siddhartha, D. Kaikho, and A. P. Madhavan for help with fieldwork. As field technicians and assistants, G.Moorthi, T. Sundarraj, T. Vanidas, and R. Rajesh provided invaluable assistance in the field, besides A. Sathish Kumar who assisted with logistics. We thank P. Pavithra for help with data entry and compilation. A special thanks to Venky Muthiah, Parry Agro Industries Limited, and several managers including Arunkumar Menon, Murali Padikkal, Mahesh Nair, Oliver Praveenkumar, K. Krishnakumar, A. K. Pradeep Kumar, G. P. Alexander, M. Ravi, and S. Marimuthu for their support and partnership.

## Data Availability Statement

The dataset and code related to this manuscript are available on Zenodo (Murali *et al*. 2023). The dataset will be made public upon acceptance of the manuscript for publication.

